# Gene family expansions underpin context-dependency of the oldest mycorrhizal symbiosis

**DOI:** 10.1101/2024.04.12.588857

**Authors:** Damian J. Hernandez, Gwendolyn B. Pohlmann, Michelle E. Afkhami

## Abstract

As environments worldwide change at unprecedented rates during the Anthropocene, understanding context-dependency – how species regulate interactions to match changing environments – is crucial. However, generalizable molecular mechanisms underpinning context-dependency remain elusive. Combining comparative genomics across 42 angiosperms with transcriptomics, genome-wide association mapping, and gene duplication origin analyses, we show for the first time that gene family expansions undergird context-dependent regulation of species interactions. Gene families expanded in mycorrhizal fungi-associating plants display up to 200% more context-dependent gene expression and double the genetic variation associated with mycorrhizal benefits to plant fitness. Moreover, we discover these gene family expansions arise primarily from tandem duplications with >2-times more tandem duplications genome-wide, indicating gene family expansions continuously supply genetic variation allowing fine-tuning of context-dependency in species interactions throughout plant evolution.

**One-Sentence Summary:** Gene family expansions arising from tandem duplications underpin genetic regulation and fitness effects of context-dependency

## Main Text

Context-dependency is one of the oldest and most pervasive concepts in biology, affecting organisms across the entire tree of life. For instance, all interactions among species change depending on environmental context (*1, 2*) and this context-dependency determines how organisms thrive in their habitats (*2, 3*), how communities assemble (*4, 5*), and how ecosystems maintain biodiversity and crucial services (*6, 7*). While context-dependency of species interactions is ubiquitous and well-documented, the genomic and molecular mechanisms allowing species to integrate environmental cues and modulate their responses to match changing environmental conditions remain largely unknown. Identifying the molecular bases of context-dependency will be particularly important in the Anthropocene as humans drive environments to new extremes through climate change, pollution, and other escalating stress. Without a clearer mechanistic understanding of context-dependency, we are running into new environmental extremes without knowing which species interactions will be resilient to intensifying anthropogenic forces. Here, we discover an essential molecular mechanism underlying context-dependency of species interactions by demonstrating gene family expansions underpin context-dependent gene expression regulating interactions and genotypic differences in fitness benefits resulting from these interactions. Further, we identify widespread tandem duplications as a ubiquitous evolutionary mechanism generating these important gene family expansions across angiosperms, linking context-dependency of species interactions to their molecular, genomic, and evolutionary mechanisms for the first time.

In the real world, organisms navigate complex environments in which conditions frequently change. Success of organisms living under changing conditions often requires the flexibility to efficiently finetune their physiological and molecular responses, which is inherently challenging and likely requires complex regulatory mechanisms. In this study, we demonstrate that the creation/maintenance of large gene families – *i*.*e*., related genes descending from a common ancestor – is an essential evolutionary mechanism underpinning how organisms finetune species interactions in response to environmental cues (Fig. 1). Large gene families offer opportunities for flexibility in organisms’ responses to their environment by creating more points of genetic regulation (i.e., complexity) which allows detailed control of expression of genes with biochemically similar functions under unique combinations of environmental conditions (e.g., *combinatorial coding* in Fig. 1; *8, 9*). The genetic complexity created through gene family expansions could be a fundamental mechanism underlying the gene expression plasticity species need to respond to different environmental conditions. As gene families expand through processes like whole-genome and tandem duplications, new genes can then be selected for new functions (e.g., *neofunctionalization* in Fig. 1; *10, 11*). For example, in yeast, gene duplication often leads to one copy maintaining its original function and the other developing a new stress-related function (*10*). While it is well-established that gene family expansions are critical for creating genetic complexity underlying novel structures and body plans (*11*–*13*) or adaptation to new environments (*14, 15*), it remains unknown if gene family expansions are important for integrating environmental cues in the context-dependency of species interactions.

**Fig. 1.**
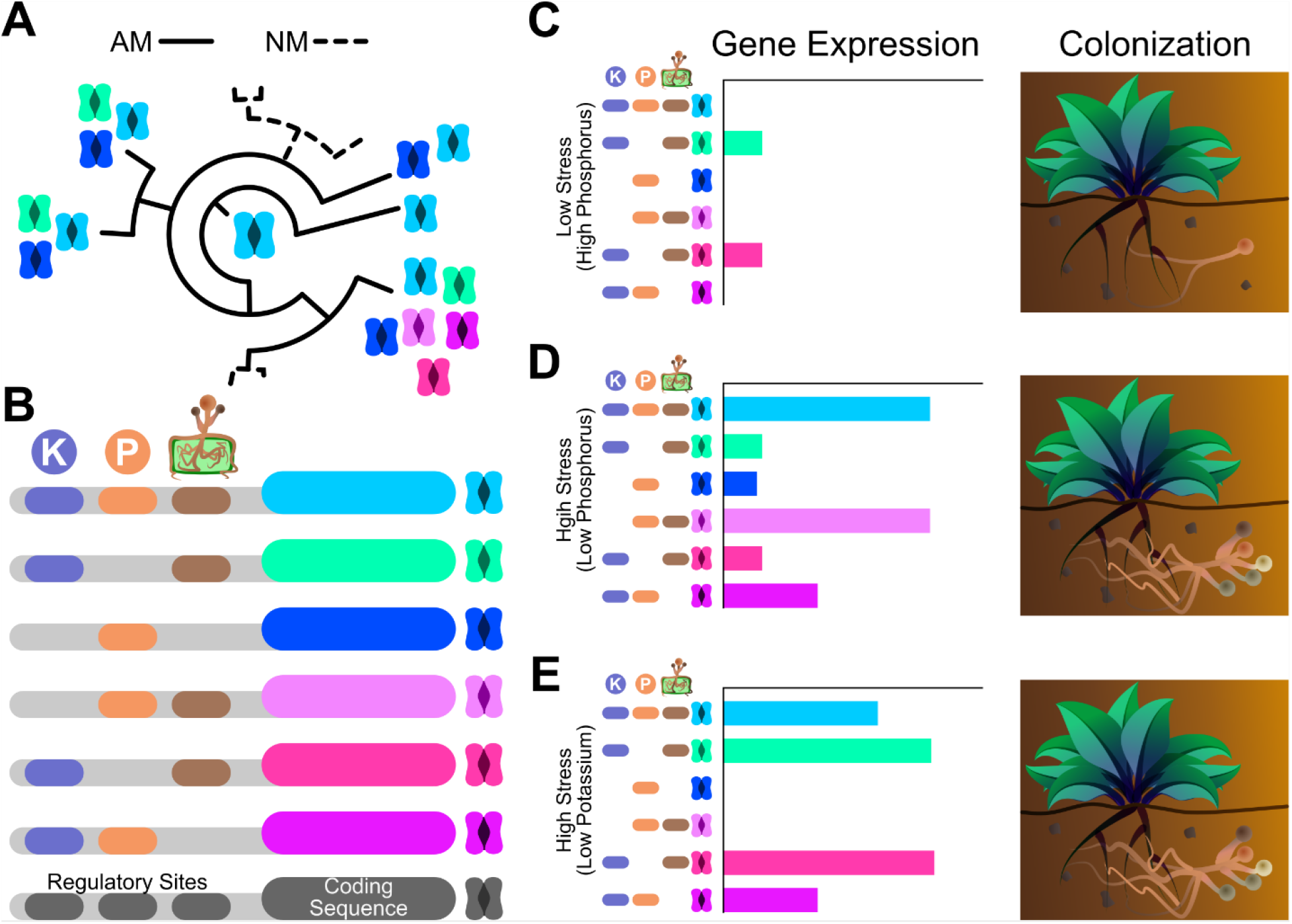
Conceptual framework for how gene families expanded in AM fungi-associating plants could allow integration of multiple environmental cues. (**A**) An original ancestor gene (central light blue) that regulates AM symbiosis could evolve into new variants (different colors), resulting in expanded gene families. In contrast, genes can be lost in plants that do not maintain AM symbiosis (dashed line lineages; NM: non-mycorrhizal) because there is no selective pressure to keep AM-relevant genes. (**B**) The evolution of new genes (colors other than light blue) through expansions can allow for neofunctionalization/subfunctionalization through combinatorial coding in which different genes in the family respond to unique combinations of environmental factors like phosphorus availability, potassium availability, and mycorrhizal presence. Note that gene family expansions may also integrate environmental cues into regulation of context-dependent interactions through other pathways not shown here, such as the gain/loss of regulatory target sites on protein sequences. (**C**-**E**) Organisms exist in complex and dynamic environments where gene expression changes based on the conditions organisms experience. In diagrams (C-E), we represent the gene family in (B) whose expression promotes AM fungal colonization but whose genes are differentially expressed in different contexts. The total expression of this gene family determines colonization where downregulation of the whole family inhibits colonization (C) and upregulation promotes it (D, E). Thus, even though the patterns of expression may be different, if the total expression of the gene family remains the same, the outcome for the interaction is the same (e.g., upregulation of different sets of genes within the family in D and E both lead to colonization). This outcome of similar total expression but different patterns of expression among genes within families may arise from combinatorial coding in the regulatory sites in which some genes respond only to phosphorus and mycorrhizae (pink gene), only potassium and mycorrhizae (cyan and hot pink genes), or a balance between all three (light blue gene).

Additionally, determining the evolutionary origins of gene family expansions underpinning context dependency is important for understanding whether the regulation of context-dependency is dramatically reengineered a few times throughout evolution (e.g., widespread, simultaneous selection across the genome in whole-genome duplications; *15*–*19*) or if the regulation of context dependency is gradually modified over time (e.g., selection on a single gene following tandem duplications; *20*–*22*). For example, a whole-genome duplication in legumes reengineered entire symbiotic pathways and a whole organ devoted to rhizobial mutualism (*11, 23*). At the other extreme, single-gene duplications (e.g., tandem duplications) limit effects to individual genes and often impact only one step of a pathway or a few downstream processes (*20*–*22*). This form of gene duplication can expand the functional repertoire of a specific class of gene without disrupting broader, essential pathways. These two different evolutionary origins of gene family expansions have other important consequences for macro- and micro-evolutionary trajectories. Whole-genome duplications, which are rare compared to tandem duplications (*17*), increase the probability of speciation (*24*–*26*), and thus also likely reduce the flow of newly duplicated genes into the original species’ populations (i.e., new gene variants flow into new species rather than populations of the original species).

Alternatively, since tandem duplications cause speciation less frequently, they can provide a continuous source of genetic novelty within species (*17, 27*). In short, discovering the evolutionary origin of duplicated gene families is essential for predicting the consequences to species’ ecology, evolution, and the molecular restructuring of context-dependent processes.

Here, we use one of the most ubiquitous and ecologically-important symbioses on Earth – the interaction between arbuscular mycorrhizal (AM) fungi and plants – to test whether expanded gene families play outsized roles in context-dependency of beneficial interactions and how these gene family expansions evolve. Up to 80% of all land plants form intracellular associations with AM fungi in which plants provide fungi with carbon in exchange for difficult-to-acquire nutrients like phosphorus (*28*). AM symbiosis is also an ancestral trait of most land plants (*29, 30*) extending back ∼450 million years ago when plants were just colonizing land (*31, 32*). Consequently, AM symbiosis is an excellent model to understand the evolution of mechanisms regulating context dependency because it has a long, shared evolutionary history across nearly the entire plant kingdom but also has replicated, independent losses. Together, this makes it possible to identify gene families regulating AM symbiosis with impacts on gene expression and mycorrhizal benefits to plant fitness and determine if gene families regulating AM symbiosis expand through the same evolutionary mechanism across the plant kingdom. We combine comparative genomics across 42 angiosperms with transcriptomics, genome-wide association study (GWAS) mapping, and gene duplication origin analysis to: 1) identify expanded gene families in mycorrhizal-associating plants, 2) demonstrate that gene family expansions are integral to context-dependent gene expression and mycorrhizal benefits to plant fitness and 3) discover that gene families expanded in AM-associating plants primarily grow through tandem duplications. Taken together, our work establishes a molecular eco-evolutionary principle for context-dependency in which evolution of molecular complexity is essential for regulating species interactions in different contexts and highlights how gene family expansions and widespread local-scale duplications are integral to evolving that complexity.

## Results

To identify gene families expanded in AM plants, we used comparative genomic analyses across 42 plant species (32 AM, 10 non-mycorrhizal) to determine which gene families were significantly larger in plants that can symbiose with AM fungi (Methods; *29, 30*). We identified 429 gene families (1.5% of 28,756 families) expanded in AM plants (Fig. 2) and found several lines of evidence supporting the role these AM-expanded gene families play in AM symbiosis.

**Fig. 2.**
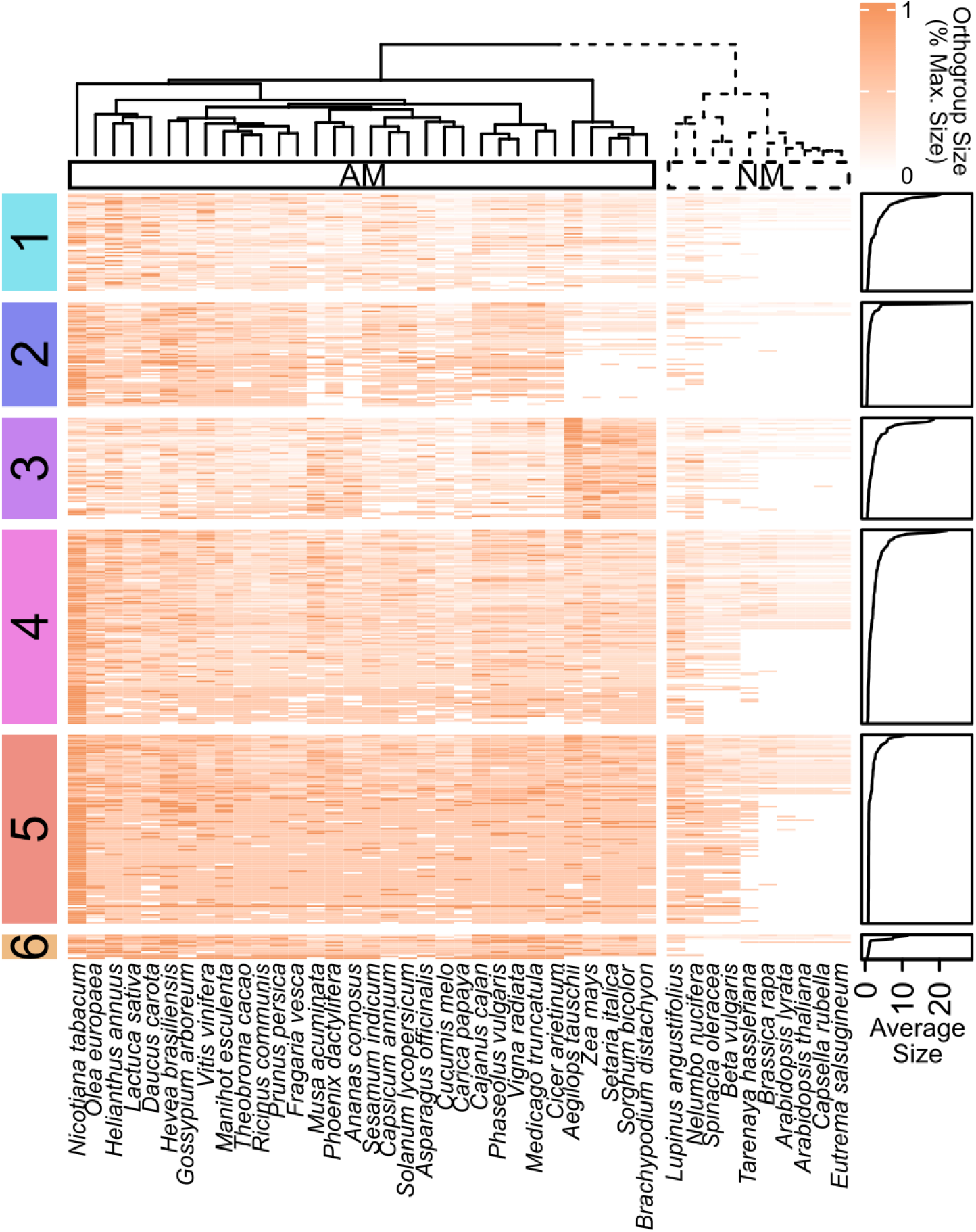
Comparative genomics identified 429 AM-expanded gene families as candidates for context-dependent regulation of the widespread symbiosis between mycorrhizal fungi and plants. Our comparative genomic analyses identified 429 gene families that were significantly larger in AM plants across 32 arbuscular mycorrhizal (AM, left) and 10 non-mycorrhizal (NM, right) plant species (Mann-Whitney U-Tests, Benjamini-Hochberg correction). We find interesting co-expansion patterns of gene families across AM plants and lineage-specific patterns in grasses (see Supplementary Text for details). For example, Cluster 1 is highly enriched in genes localized to membranes and arbuscules such as the EIX2 fungal receptors and Vapyrin-like proteins that may organize arbuscule development. Each row represents a gene family with darker colors indicating larger gene families. Plant species (columns) are clustered within the heatmap using unsupervised hierarchical clustering (Ward’s D2 linkage, Euclidean distances) on gene family sizes, which neatly split AM and NM plants into two distinct groups. We used k-means clustering on gene families (rows) to identify six groups (colored labels on left). To allow for comparison of gene families across rows, values within each cell were normalized across the row (values between 0 and 1).

First, 70% of genes in the Common Symbiosis Pathway (a pathway required for establishing AM symbiosis; *33*) are in AM-expanded gene families, which is 47-times greater than the null expectation of 1.5% of all gene families (p < 0.0001, binomial test; fig. S1a). In fact, only the three nucleoporins *SEH1, NUP85*, and *NUP133* were not in AM-expanded gene families, suggesting that the nuclear transport organized by these proteins may be essential for AM establishment but not unique to it, unlike processes organized by genes mediating nuclear calcium spiking (*CASTOR* and *DMI1/POLLUX*), inducing hyphopodia formation (*RAM2*), and reprogramming transcription (*DMI3/CCamK, IPD3/CYCLOPS, RAM1*, and *NSP1*), which are essential and unique to AM symbiosis or recently co-opted for other symbioses. Second, we find genes within AM-expanded gene families are 63% more likely to be differentially expressed than the rest of the genome when plants are grown with AM fungi compared to when they are not (p < 0.0001, binomial test; fig. S1b; *34*), thus supporting expression of genes in AM-expanded gene families as an important regulatory target for plants during AM symbiosis. Our analyses also revealed interesting co-expanding groups of gene families, providing insight into co-regulated processes in AM symbiosis across angiosperms and modifications to symbiosis in grasses (see Supplementary Text, Supplementary File 1). Taken together, our comparative genomics analysis identifies gene families likely functioning in AM symbiosis across plants whose last common ancestor was ∼160-200 million years ago (*35*).

We then asked whether larger AM-expanded gene families are hotspots of context-dependent regulation and mycorrhizal benefits to plant fitness. If gene family expansions underpin context-dependent regulation of AM symbiosis, plants should disproportionately regulate genes in larger AM-expanded gene families in response to the combination of AM symbiosis and environmental stress rather than these two factors separately (i.e., significant interactive effects of mycorrhizal fungi and environment indicate context-dependent gene expression). Also, if larger AM-expanded gene families are important for context-dependent regulation of AM symbiosis and, consequently, fitness outcomes in associating plants, then larger gene families should contain more genetic variation associated with mycorrhizal-derived fitness benefits than the rest of the genome. Therefore, we 1) conduct a context-dependent expression analysis to determine if plants disproportionately regulate expression of genes in larger gene families during AM symbiosis in stressful conditions and 2) perform a GWAS to determine if larger gene families are hotspots of genotypic differences in mycorrhizal-derived benefits to plant fitness.

First, we found that larger AM-expanded gene families had up to 200% greater context-dependent expression under soil chemical stress than the rest of the plant genome (Fig. 3a), indicating that gene expression plasticity facilitated by expanded gene families provides an important genomic basis for context-dependent regulation. We analyzed RNA-sequencing datasets of plants grown in factorial experiments with/without AM fungi and with/without a stressor – either low phosphorus, low potassium, high salinity, or drought (*36*–*39*). We then determined if larger AM-expanded gene families had significantly higher proportions of genes whose expression is context-dependent. To do this, we quantified the relationship between gene family size and proportion of genes in AM-expanded gene families with significant context-dependent expression compared to the same relationship in 10,000 random subsamples of all gene families. We use proportion (instead of total number) of differentially-expressed genes to avoid a size bias (see Methods). We found plants disproportionately target larger AM-expanded gene families for context-dependent regulation under soil chemical stress, with 200%, 86%, and 41% greater context-dependent expression of these gene families than expected by chance in response to low phosphorus, low potassium, and high salinity stress, respectively (p = 0.0006, p = 0.0054, and p = 0.0297; Fig. 3a), indicating the gene expression plasticity facilitated by gene family expansions is important for context-dependent regulation under soil chemical stress.

**Fig. 3.**
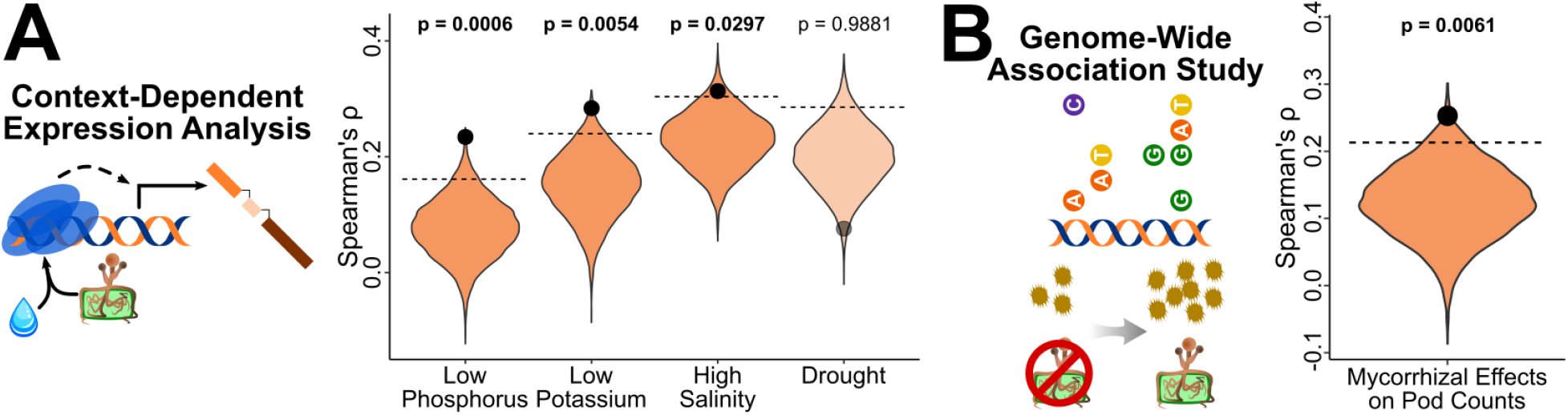
Larger AM-expanded gene families are hotspots of context-dependent gene regulation and mycorrhizal benefits to host plant fitness. (**A**) <u>Our transcriptomic analyses discovered that larger AM-expanded gene families have 40-200% more context-dependent expression in response to soil chemical stress and mycorrhizal presence than expected by chance.</u> We quantified the strength of the relationship between each gene family’s size and the proportion of genes in each family that were context-dependently expressed (Spearman’s Correlation Coefficients; black dots) and compared these observed relationships to the null expectation generated by permutational analysis with 10,000 random subsamples of all gene families (violins). (**B**) <u>Our genome-wide association study (GWAS) also identified larger AM-expanded gene families as genetic hotspots for mycorrhizal benefits to plant fitness, harboring ∼100% more associated SNPs than the rest of the genome</u>. To determine if larger AM-expanded gene families contain more genetic variation underlying naturally-occurring variation in plant fitness benefits from AM fungi, we determined the number of SNPs associated with increases (or decreases) in pod counts in 212 *M. truncatula* genotypes grown with versus without AM fungi using the GWAS. We then quantified the relationship between the normalized SNP density of each gene family (i.e., number of significantly associated SNPs divided by the entire sequence length of each gene family which accounts for variation in the number of genes and gene lengths in different families) and related normalized SNP density to gene family size using a Spearman’s correlation test (observed coefficient displayed as black dot). To determine if those observed relationships were significant, we compared the observed coefficient to that from 10,000 random subsamples of all gene families (violin). In (A) and (B), the dashed lines represent the 95% significance cut-offs; actual coefficients (black dots) above these lines indicate a significantly stronger relationship between gene family size and context-dependent expression or between gene family size and genetic variation than expected by chance. Faded points and violins represent non-significant relationships.

Interestingly, under drought, larger gene families do not contain more context-dependent expression (observed Spearman’s ρ: 0.076; mean random ρ: 0.199; p = 0.9881; Fig. 3a) but may actually contain less (p = 0.0119; 61% less than random on average). This dichotomy between soil chemical stressors and drought potentially points to two distinct molecular syndromes driving the ecological strategies plants use to regulate AM symbiosis during persistent stress lasting years/decades versus seasonal/episodic stress (see Supplementary Text). In both cases, our transcriptomic analyses demonstrate that gene family size is an essential component of how plants regulate gene expression when integrating environmental context into AM symbiosis.

Second, larger AM-expanded gene families harbor 2-times more genetic variation associated with mycorrhizal benefits to plant fitness than the rest of the plant genome, highlighting how gene family expansions can underpin significant intraspecific variation in mycorrhizal benefits. Motivated by our finding that plants disproportionately target larger gene families for regulating gene expression during AM symbiosis under soil chemical stress, we evaluated whether these AM-expanded gene families disproportionately impact mycorrhizal benefits to plant fitness by conducting a GWAS across 212 *Medicago truncatula* genotypes (*40, 41*). The GWAS identified genetic variation – SNPs, specifically – linked to mycorrhizal effects on the key plant fitness metric of seed pod counts (*42*). We used permutational tests to determine if larger AM-expanded gene families contained more fitness-associated SNPs than expected (compared to 10,000 random subsamples of the entire genome). Larger AM-expanded gene families contained approximately double the genetic variation underpinning mycorrhizal effects on fitness than expected by chance (observed Spearman’s ⍴ = 0.252, mean random ⍴ = 0.125 ± 0.001; p = 0.0061; Fig. 3b), indicating these gene families are hotspots for genetic variation underlying mycorrhizal effects on plant fitness. This emphasizes that in addition to being focal points for soil chemistry-driven regulation, larger AM-expanded gene families are an important source of the genetic variation determining mycorrhizal benefits.

Our result that gene family expansion is an important genomic mechanism for context-dependent regulation and benefits of AM symbiosis raises the question: how are these gene families expanding? Whether expansions occur through massive duplication events like whole-genome duplications or through smaller, single-gene duplications can have profound consequences for how frequently new genes are created (*17*) and the extent that molecular pathways are reengineered (*18, 43*). Consequently, we sought to determine which duplications were the main drivers of growth for AM-expanded gene families.

We find that AM-expanded gene families grow >2-times as often through tandem duplications (duplication of genes immediately adjacent to each other on a chromosome) as the rest of the genome across all plant species tested (Fig. 4), indicating that tandem duplication is the primary evolutionary driver of expanding gene families underpinning context-dependency of this symbiosis. We conducted a gene duplication origin analysis using a customized *MCScanX* pipeline (DOI: 10.5281/zenodo.10934057; *44*) and quantified the frequency of different types of duplications in AM-expanded gene families across 24 plant species with high-quality chromosome scaffolds (results are also consistent across all 32 AM-associating species; fig. S5). We then compared those observed frequencies with 10,000 random subsamples of gene families from each species’ genome (Methods). In all AM-associating plant species (24/24, FDR < 0.05), genes in AM-expanded families disproportionately originate from tandem duplications (2.28 ± 0.08 times more common than random; Fig. 4b,c). This is further supported by modest enrichment of proximal duplications (duplications with 1-20 genes between duplicates; Methods) in 42% of plant species (10/24 significant and all species except *Zea mays* show the same trend; Fig. 4b,d) because many proximal duplications are older tandem duplications that have been interrupted by events like gene insertion (*17, 44*). Dispersed and whole-genome/segmental duplications are negligible drivers of growth in AM-expanded gene families (0% and 8.33% of species, respectively; Fig. 4e,f). Taken together, these results demonstrate that tandem duplication is the primary evolutionary source of AM-expanded gene families, highlighting that local, small-scale duplications have profound impacts on context-dependent regulation across angiosperms.

**Fig. 4.**
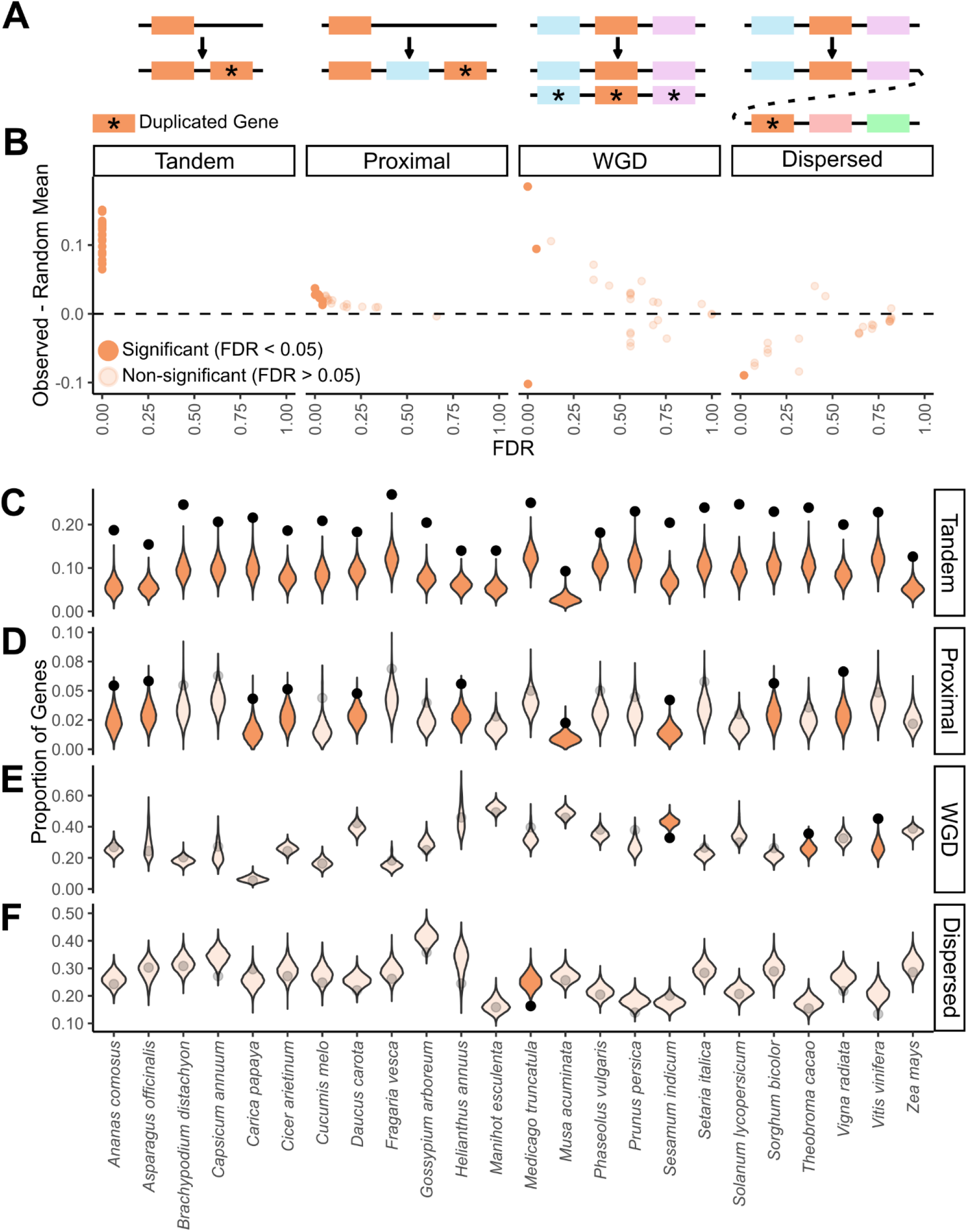
Tandem duplications are the main driver of growth in AM-expanded gene families across all angiosperm species examined. (**A**) <u>Cartoon schematic of four duplication classes evaluated – tandem, proximal, segmental/whole-genome duplication (WGD), and dispersed duplications –</u> with focal genes in orange and duplicates genes marked by asterisks. **(B)** <u>Frequency of different duplication events in AM-expanded gene families compared to the whole genome across 24 angiosperms</u>. Each point represents one angiosperm species. Y-axis displays the difference between the actual frequency of that type of duplication in AM-expanded gene families and the null expectation (averages from 10,000 randomizations; displayed as violin distributions in C-F). X-axis shows the Benjamini-Hochberg-corrected p-values (FDR) of each comparison between the actual frequency in AM-expanded gene families and the frequencies in the random subsamples. Significant points above the dashed line (dashed line represents no difference between observed and random mean) represent significant enrichment of that duplication type in that species and significant points below the dashed line represent depletion. **(C-F)** <u>Violin plots showing which type of duplications occur significantly more (or less) often in AM-expanded gene families than expected by chance for each of the 24 plant species.</u> Black points represent the proportion of genes in AM-expanded gene families originating from tandem (C), proximal (D), whole-genome/segmental (E), and dispersed (F) duplications found in our analyses. Violin distributions display the null expectations (i.e., distribution of genes that were classified as each duplication type in 10,000 random subsamples of all gene families) for all 24 tested species. Unfaded points and violins are significant relationships (FDR < 0.05; observed proportion of duplications outside of 95% of the null distribution) and faded points and violins are non-significant.

## Discussion

Our study demonstrates for the first time that (1) gene family expansions are an important mechanism through which species build molecular complexity needed to regulate interactions in changing environmental conditions, and (2) tandem duplications are the main evolutionary mechanism facilitating these molecular and genomic mechanisms (gene expression plasticity and gene family expansion, respectively) underlying context dependency. Using the widespread and ecologically-important symbiosis between plants and arbuscular mycorrhizal fungi as a model, we found that expanded gene families are hotspots for both context-dependent genetic regulation and genetic variation underpinning organismal fitness. Surprisingly, expanded gene families experienced up to 200% more context-dependent changes in molecular regulation of the symbiosis in response to soil chemistry changes and twice as many fitness-associated SNPs as the rest of the genome, emphasizing the importance of this mechanism for providing molecular plasticity required for context dependency and modulating fitness benefits of symbiosis. Our results also highlight how influential micro-evolutionary processes – widespread tandem duplication – can shape how organisms regulate their interactions. In fact, expanded gene families regulating AM symbiosis primarily grew through tandem duplications across all 32 tested plant species, with >2-times more tandem duplications in AM-expanded gene families than the rest of the genome. Below, we discuss 1) how evolutionary processes driving gene family expansions shape species’ ecologies in changing environments and 2) how integrating molecular biology into ecological frameworks generate valuable mechanistic understanding of complex ecological processes.

Our research highlights tandem duplications as a key source of genetic variation for context-dependent regulation of symbioses. Because tandem duplications continuously occur throughout plant evolution (*17*) without the high likelihood of speciation resulting from more disruptive processes like whole-genome duplications (*24*–*26, 45*), tandem duplications can provide new genes that spread through species’ populations and increase the functional repertoire for regulating context-dependency *within* species (i.e., microevolution). This contrasts with whole-genome duplications which are also important for creating evolutionary novelty but can generate macroevolutionary changes like speciation and radiation (e.g., as in legumes, angiosperms, etc.; *18, 26, 45*) that can reduce the flow of newly duplicated genes back into the original species’ populations. Consequently, tandem duplications may be important for generating variation in gene copy numbers within a population that can be selected on by natural selection to shape the fitness consequences of context-dependency. Our results support this evolutionary mechanism by discovering enrichment of context-dependent gene regulation and fitness-associated variation in larger gene families. In short, our work highlights the importance of tandem duplications as a main driving force in the context-dependent regulation of AM symbiosis. We hypothesize tandem duplications and gene family expansions will be a common mechanism for context-dependent regulation of species interactions across the tree of life and advocate for future work across a wide range of species interactions.

Our work also illustrates how integrating molecular biology with ecological theory can meaningfully expand our understanding of context-dependent mechanisms and their biological consequences. Historically, researchers of context-dependency have built value-based models (e.g., Biological Market Theory, Stress Gradient Hypothesis, etc.; *46, 47*) in which each species provides a resource whose value shifts based on the environmental context (e.g., mycorrhizal-derived phosphorus has more value in low-phosphorus soils than in high-phosphorus soils).

These value-based models provide easy-to-understand, ecologically-meaningful reasons why interactions change. Genome-wide approaches, like we use here, provide an important complementary perspective by generating holistic insight into how species determine the value of resources in different environments. For example, by using comparative genomics, we identified unique gene family co-expansion in specific plant lineages that highlight likely changes in how species value certain resources. For instance, several co-expanding gene families involved in starch/carbon metabolism are highly reduced in grasses when compared to other plants (e.g., OG0013009 involved in glycogen metabolism and OG0013006 involved in sucrose catabolism; Supplementary File 1), suggesting the value of carbon in AM symbiosis is altered in grasses and emphasizing the importance of generating grass-specific models of carbon-phosphorus trade in AM symbiosis rather than inferring trade dynamics from symbioses with other plant lineages. Further, our study highlights a pertinent frontier at the intersection of mechanistic understanding of context-dependency and global change biology in which gene family expansions can create genetic variation within species that is used to regulate their interactions and potentially better navigate new, more stressful extremes in the Anthropocene.

However, our study showed the importance of expanded gene families as a mechanism for modulating beneficial species interactions may depend on the type of stressor with different mechanisms for persistent and seasonal/episodic stresses, such that the complexity generated by gene family expansions may be more beneficial under persistent stressors like low nutrient availability but disadvantageous under episodic stressors like drought (Supplementary Text).

Future work exploring how molecular complexity relates to the temporal characteristics of stressors would be useful for linking genomic data to the susceptibility of different species to their new environmental conditions in the Anthropocene (e.g., more intense/frequent droughts, warmer temperatures, etc.). Overall, we predict a flourishing of innovative molecular perspectives applied to context-dependency will be possible as massive collaborative efforts to expand molecular databases come online (e.g., 10000 Plant Genome Project for plants, i5k for insects, Refuge 2022 Consortium for corals, etc.; *48*–*50*), thus highlighting how the current state of the field is at the cusp of linking molecular mechanisms to large-scale, ecological consequences.

In summary, gene family expansions are an essential mechanism for developing complexity needed to regulate context-dependency of species interactions. We demonstrate this in the widespread mycorrhizal symbiosis in plants with ∼200 million years of shared evolutionary history. Specifically, we show how plants disproportionately regulate the largest AM-expanded gene families and that these gene families contain almost double the amount of genetic variation associated with mycorrhizal-derived plant fitness. We also find a ubiquitous evolutionary mechanism through which gene family expansions occur in this interaction, highlighting how small, local duplications may be essential to microevolution of context-dependent regulation of interactions. Our work demonstrates that complexity is not obfuscating the molecular mechanisms underlying context-dependency but is actually an important tool underlying context-dependent regulation of interactions. Consequently, we should embrace the complexity of context-dependency and explore where and when complexity originates as species interactions evolve. Discovering how complexity is generated will prove integral to understanding not only under which conditions an interaction will be beneficial or harmful but also how processes occurring at a molecular level cascade to shape interactions between entire species.

## Acknowledgements

We would like to thank Drs. Alex CC Wilson (University of Miami), Julia Dallman (University of Miami), Stefan Wuchty (University of Miami), and Javier del Campo (University of Miami/Universitat Pompeu Fabra) for their advice during this project and feedback on this manuscript. We would also like to thank the Christopher A Searcy lab (University of Miami), Dr. John Stinchcombe (University of Toronto), and Dr. Vasilis Kokkoris (Vrije Universiteit Amsterdam) for their suggestions in the writing of this manuscript.

## Funding

University of Miami – Maytag Fellowship (DJH)

University of Miami – Dean’s Summer Research Fellowship (DJH)

University of Miami – Dean’s Dissertation Fellowship (DJH)

USDA NIFA Predoctoral Fellowship (2022-67011-36456) (DJH)

University of Miami – College of Arts and Sciences Summer Research program (GBP)

NSF DEB-1922521 (MEA)

NSF DEB-2030060 (MEA)

NSF DBI-2305481 (DJH)

## Author contributions

Conceptualization: DJH, MEA Methodology: DJH, GBP Visualization: DJH Funding acquisition: DJH, GBP Supervision: DJH, MEA Writing: DJH, GBP, MEA

## Data and Materials Availability

Raw data are publicly available and are linked to the public databases from which they were retrieved in a Zenodo repository (DOI: 10.5281/zenodo.10934057). Scripts to analyze the data and processed data for/from statistical analyses are available in Zenodo. Metadata and processed data used in statistical analyses are also available in Supplementary File 1.

