## Supplementary Materials for "Gene family expansions underpin context-dependency of the oldest mycorrhizal symbiosis"

Materials and Methods

All analyses were performed in *R* (v 4.1.2) with the packages listed in Supplementary File 1 unless otherwise indicated. All original data are publicly available and can be found as indicated in Supplementary File 1 or in our Zenodo repository (DOI: 10.5281/zenodo.10934057). Computational scripts, processed data, and analyses are fully available in Zenodo. Processed data are also available in Supplementary File 1.

Identifying gene families across 42 plant species

For our comparative genomic analysis of gene family expansions, we first identified gene families across 42 angiosperms. We included plant species with 1) well-documented mycorrhizal status, 2) importance as agricultural/research plants, and 3) well-characterized genome annotations. We determined mycorrhizal status (whether or not it can symbiose with arbuscular mycorrhizal fungi) based on agreement between *MycoDB* (*51*), the widely used, global database of plant mycorrhizal fungi-plant interactions, and Maherali et al (*30*), which phylogenetically reconstructed mycorrhizal status across seed plants (i.e., ~380 million years of plant evolution). We restricted our analyses to RefSeq annotations because all genomes in RefSeq have been processed through NCBI’s Eukaryotic Genome Annotation Pipeline which uses the same gene prediction algorithms and validates gene models with evidence from RNA-sequencing datasets thus ensuring all considered genomes experienced a similar annotation process. We also focused on plant species that do not associate with ectomycorrhizal fungi because plants repurposed some genes involved in arbuscular mycorrhizal (AM) symbiosis for their new symbiosis with ectomycorrhizal fungi (*52*–*54*). Including ectomycorrhizal plants in our analyses would have identified gene families involved in an interaction unrelated to AM symbiosis and thus outside the scope of our question. Based on these criteria, we identified 42 plant species that could be used in our study: 32 plants that associate with arbuscular mycorrhizal fungi (AM) and 10 that do not (non-mycorrhizal, NM; Supplementary File 1) and downloaded all protein-coding sequences for each species from NCBI’s RefSeq database. For input into the algorithm grouping genes/proteins into gene families (i.e., *OrthoFinder*), we grouped proteins by their gene identifier (GI) to associate different protein isoforms with their respective gene. We then selected the longest protein isoform of each gene for input into *OrthoFinder* as suggested by the program writers. This isoform selection ensures that gene counts within families are not inflated by isoforms.

We then grouped genes into “orthogroups” using the *OrthoFinder* algorithm (v2.2.6; *55*, *56*) with default parameters. Orthogroups are sets of genes descending from a single gene in the common ancestor and this can encompass both orthologs and paralogs. In this manuscript, we refer to orthogroups hereafter as gene families to ensure consistent and accessible terminology. *OrthoFinder* predicts gene families by first performing reciprocal *DIAMOND* (a resource-efficient and optimized version of *BLAST*; *55*–*57*) searches on all proteins in a proteome database with a gene length correction to account for statistical biases in alignment scores from gene lengths. After identifying reciprocal best hits, proteins are organized into networks. These networks are then subdivided into gene families using a Markov Clustering Algorithm. To further finetune the gene families, *OrthoFinder* then constructs a species tree from the gene trees and uses the newly constructed species tree to identify gene duplication events. *OrthoFinder* uses these gene duplication data to improve gene family identification by resolving gene trees and correcting gene families within the context of duplication events (*56*).

Identifying gene families expanded in arbuscular mycorrhizal plants

To identify which gene families are expanded in AM plants, we compared the sizes of gene families between AM and NM plants (i.e. between plants that associate with arbuscular mycorrhizal fungi and non-mycorrhizal plants) using Mann-Whitney U-tests for each gene family and then corrected for multiple comparisons with a Benjamini-Hochberg correction. Gene families that were significantly larger in AM plants compared to NM plants were considered putative AM-expanded gene families. The ability to associate with AM fungi is an ancestral trait for most land plants (*29*, *30*), and as a result, gene families that are smaller in NM plants are likely involved in establishing, maintaining, and/or regulating AM symbiosis because NM plants will have lost the selective pressure to maintain these families. Note that AM-expanded gene families could result from both selection for the maintenance and the enlargement of gene families over evolutionary time scales. While it is not computationally possible to differentiate between maintenance and expansion in this comparative genomics analysis, results from our gene duplication origin analyses strongly support AM-expanded gene families arising from enlargement of gene families rather than maintenance.

To verify that these putative AM-expanded gene families are involved in AM symbiosis, we validated our selection by 1) loss of known AM-symbiosis genes in NM plants and 2) determining if AM-expanded gene families are more likely to be differentially expressed in plants grown with AM fungi relative to plants not grown with AM fungi. First, we selected gene families containing genes that are part of the Common Symbiosis Pathway (CSP; *33*) and assessed if they are over-represented in the putative AM-expanded gene families. The CSP is a known pathway of genes that are essential for establishment of AM symbiosis (*33*). We used a binomial test (*binom.test*, *base R* package) to compare the observed proportion of CSP gene families that are retained/expanded in AM plants (70%) to the proportion of *all* plant gene families that are conserved in AM plants (1.5%). 2) Second, we determined if potential AM-expanded gene families are more likely to be differentially expressed in plants grown with AM fungi relative to those that are not. We analyzed RNA-sequencing data from Afkhami et al 2016 (*34*, *58*) in which *M. truncatula* was grown with and without the AM fungus *Rhizophagus irregularis* and with and without the nitrogen-fixing bacterium *Ensifer meliloti.* We compared expression in plants grown with and without the AM fungi (and without *E. meliloti*). RefSeq IDs of proteins in our gene families were matched to J. Craig Venter Institute’s *M. truncatula Jemalong A17 4.0* annotation (*59*) using the annotation table from ENSEMBL Plants (release 53). We then 1) used a binomial test (*binom.test*, *base R* package) to determine if the proportion of gene families that contain at least one differentially-expressed gene was higher than the null expectation of the proportion of all gene families containing at least one differentially-expressed gene and 2) whether genes in gene families that are expanded in AM plants are more likely to be differentially expressed than non-conserved genes using a permutational test (*independence_test*, *coin* package).

Determining if larger AM-expanded gene families have more context-dependent expression (Differential Expression Analysis)

Because we were interested in determining if large gene families are important in regulating AM symbiosis in response to different environmental stimuli, we sought to analyze RNA-sequencing datasets in which plants were grown with/without mycorrhizal fungi and with/without an environmental stressor. We searched the NCBI Gene Expression Omnibus and BioProject databases for the following search terms: “arbuscular”, “rhizophagus”, “funneliformis”, “glomus”, and “mycorrhizal”. We manually reviewed all search hits for datasets in which plants were grown in factorial experiments manipulating mycorrhizal fungi (+/- AM fungi) and environmental stimuli (+/- environmental stimuli). We also performed Google Scholar searches to identify other factorial RNA-sequencing datasets using terms such as: “AM symbiosis”, “RNA-sequencing mycorrhizae”, “AM symbiosis stress”, “*Rhizophagus* stress”, “AM symbiosis drought”, etc. We focused on RNA-Seq data sets and excluded microarray datasets because genes with high homology (e.g., genes within a gene family) may amplify within the same spot on the microarray, causing ambiguity over which gene is actually expressed.

For each AM fungi and environmental stimuli factorial dataset, we matched expressed genes to gene families. If gene count data were provided through the NCBI database or with the paper, we analyzed those tables as input for our analyses. If data were provided as raw sequencing data, we downloaded the raw fastq files from NCBI’s Sequence Read Archive then mapped genes to reference genomes and quantified their expression using the STAR aligner (*60*). Scripts used to download and process raw sequencing reads are available in the Zenodo repository associated with this manuscript (DOI: 10.5281/zenodo.10934057). To associate expressed genes with their gene family when naming did not match RefSeq (e.g., ENSEMBL gene names) or if the studied organism was not one of the 42 species originally input into *OrthoFinder*, we aligned the protein or nucleotide sequences for those organisms against the RefSeq database of a closely related species using *blastp* and *blastx*, respectively. We assigned gene family identities to the unknown proteins/genes based on the top result from the *BLAST* search. We only considered a top result meaningful if e-values were less than 0.0001. If no hit was identified for that protein/gene, then we assigned the gene family as “Other” (i.e., a likely species-specific gene).

For each dataset, we evaluated which genes showed context dependent expression in response to mycorrhizal fungi and the environmental stimuli using *DESeq2* (*61*). To do this, we generated negative binomial distribution models in *DESeq2* of gene expression for each gene with the fixed factors of AM presence/absence and environmental stress presence/absence (e.g., salinity, drought, etc.) as well as the interaction between these two terms. We then compared these “full” models against models without the interaction term using a likelihood ratio-test (FDR < 0.05) to determine if the interaction term was significant. Significant interaction terms indicate non-additive, context-dependent gene expression. The expression of these genes is context-dependently affected by both mycorrhizal fungi and the environmental stimuli in ways that cannot be explained by either mycorrhizal presence or environmental stress alone (i.e. the gene’s expression response to mycorrhizal fungi depends on the environmental conditions).

To determine if larger AM-expanded gene families are disproportionately expressed context-dependently (i.e., interaction between AM presence and stressor presence is significant), we analyzed the relationship between gene family size and the proportion of genes within that family that showed context-dependent expression using a Spearman’s correlation test. We then compared that observed correlation value against 10,000 correlations of gene family size and frequency of context-dependent expression that were calculated from 10,000 random subsamples of all gene families. For example, if a plant species had 200 AM-expanded gene families, we calculated the observed Spearman’s ⍴ between gene family size and frequency of context-dependent expression and then compared that observed relationship with those in 200 randomly selected gene families from all families 10,000 times. We calculated p-values as the number of random subsamples that had a stronger (i.e., higher ⍴) or equal relationship than the observed and then divided that number by 10,000. This permutational test allows us to determine if larger AM-expanded gene families are more likely to have context-dependent expression than would be expected by random chance. Note that by using the proportion of context-dependent genes (instead of absolute number of context-dependent genes) in this analysis, we avoid a size bias in which larger gene families would have more context-dependent expression solely because they have more genes. Using the proportion in the analysis allows us to determine if genes in larger gene families are disproportionately more likely to be context-dependently expressed.

Determining if larger AM-expanded gene families have more fitness-associated natural genetic variation (Genome-Wide Association Study)

To determine if larger AM-expanded gene families are disproportionately associated with mycorrhizal effects on plant fitness, we performed a genome-wide association study (GWAS) on 212 naturally-occurring *M. truncatula* populations and quantified how much genetic variation is associated with changes in pod counts (i.e., an important fitness metric for this annual plant) between plants grown with and without AM fungi. Genomes for the 212 plant genotypes used in this analysis were available through the *Medicago HapMap Project* (*41*). The plant fitness data (pod counts of this annual plant) were collected in a factorial experiment in which the same 212 genotypes were grown in the presence and absence of mycorrhizal fungi with replication (see Afkhami et al 2021 for details of the experiment; *42*). Using *bcftools* (*62*), we identified SNPs across the plant genotypes, then filtered to remove SNP sites with minor allele frequencies less than 5% and SNPs for which more than 10% of populations had no available sequencing data for that position. This resulted in 1,760,228 SNPs available for genomewide association mapping. We imputed missing data with the most common allele for that site (“mode” option in the *impute* function of the *LEA* package).

We conducted the GWAS using the *LEA* package (*63*), which uses latent factor mixed models (LFMMs). LFMMs are regression models that combine fixed and latent effects. The fixed effect in the context of our GWAS study is the fitness variable of interest (i.e., change in pod counts between inoculated and uninoculated plants). Latent factors are unobserved confounders that correlate with both the fixed effects and genetic data. Including latent factors in the model allows the variation associated with these background confounding effects to be partitioned out to determine if a specific SNP is associated with fitness after considering the impact of the confounding effects (e.g., after accounting for background relatedness). To determine how many latent factors to which the background confounding effects should be collapsed, we assessed how many “groupings” are present in the genetic data using an elbow plot of a principal component analysis (fig. S2), elbow plot of cross-entropy criteria from admixture coefficient analyses (fig. S3), and visual assessment of the Q-matrices from the best “run” in the admixture coefficient analyses for different numbers of ancestral populations (fig. S4). In all three cases, 6-8 latent factors demonstrated the best description of structure in the data. Our analyses were consistent for either 6, 7, or 8 latent factors, but we represent the results from analyses with 7 latent factors because this was the middle of the range and produces results representative of this range. We conducted the LFMMs with the *lfmm2* function (*LEA* package), which approximates the Markov Chain Monte Carlo algorithm of the original LFMM implementation by using the substantially more computationally efficient least-squares estimation for latent factor estimation (*63*). We corrected the p-values of each LFMM for each SNP using a Benjamini-Hochberg correction.

We then asked if larger AM-expanded gene families had more genetic variation associated with mycorrhizal effects on plant fitness (i.e., pod counts) than smaller gene families. To do this, we first concatenated the annotated gene sequences of all genes within each gene family. We then asked what percentage of that concatenated gene family had genetic variation associated with changes in pod counts between plants inoculated with AM fungi and those that were not. This approach avoided a size bias (i.e. larger gene families having more genetic variation solely because their concatenated sequences are longer) by allowing us to normalize our results and thereby make gene families comparable. We performed a Spearman’s correlation analysis to determine the strength of the relationship between gene family size and the percentage of a gene family’s genomic sequence with fitness-associated variation. We then calculated the significance of that observed relationship with a permutation test; we repeated this analysis with 10,000 random subsamples of an equal number of gene families from the whole plant genome and quantified how many of these randomized subsamples had stronger relationships between gene family size and genetic variation. This approach tells us if larger gene families expanded in AM plants are likely to have outsized roles in regulating mycorrhizal effects on plant fitness.

Identifying the duplication mechanisms driving growth of AM-expanded gene families (Duplication Origin Analysis)

To identify which duplication mechanisms are driving growth in AM-expanded gene families, we performed a gene duplication origin analysis in AM-expanded gene families across 24 species whose genome annotations are complete or nearly complete (i.e., they have chromosome-level scaffolds with few other scaffolds remaining). We based our gene duplication origin analyses on the implementation of the *MCScanX* package (*44*).

We identified four different types of duplications based on how they are classified in *MCScanX*: 1) tandem duplications – genes within a family that immediately occur next to each other on a chromosome/scaffold; 2) proximal duplications – genes within a gene family that are separated by 1-20 genes; 3) dispersed duplications – genes within a gene family that are neither proximal, tandem, nor part of a syntenic block; and 4) WGD or segmental duplications – genes within a gene family that belong to a syntenic block within a species’ genome as identified by the *MCScanX_h* function of the *MCScanX* package. Genes that have no other gene family members are classified as singletons. If genes meet criteria for multiple duplication types, duplication type was assigned based on the following priority (as is done in the original *duplicate_gene_classifier* function of *MCScanX*; *44*): WGD > tandem > proximal > dispersed > singleton. We developed a custom implementation of the *duplicate_gene_classifier* function because the original *duplicate_gene_classifier* function in *MCScanX* determines gene duplicates through pairwise *DIAMOND* alignments due to using the *MCScanX* function to identify WGD/segmental duplications. However, this strategy of using pairwise alignments is less accurate than using *OrthoFinder* to identify gene duplicates/families. As a result, we recreated the *duplicate_gene_classifier* logic in *R* for tandem, proximal, dispersed, and singleton classifications to take advantage of the greater accuracy from *Orthofinder* for identifying gene duplicates/families. To identify WGD/segmental duplications, we used the *MCScanX_h* function (intraspecific-only; `-b 1` keyword argument) which can use the homology file from Orthofinder to identify WGD/segmental duplications (unlike the *MCScanX* function which can only use less accurate pairwise *DIAMOND* alignments). In short, we recreated the *duplicate_gene_classifier* logic in *R* to take advantage of the greater accuracy of our *Orthofinder* gene family groupings.

To determine if AM-expanded gene families disproportionately arise from any one type of duplication event, we calculated the observed proportion of genes in AM-expanded gene families that belonged to each type of duplication for each species. We compared those observed proportions to 10,000 random subsamples of all gene families in each species (number of randomly subsampled gene families is equal to the number of AM-expanded gene families in that species). To calculate significance, we determined how many subsamples had proportions whose absolute difference from the random mean were greater than the absolute difference between the random mean and the observed proportion in AM-expanded families. That number was divided by 10,000 (i.e., the number of random subsamples) to calculate the p-value. We then used Benjamini-Hochberg correction for the 24 permutational tests (i.e., the number of species used in the analysis) within a type of duplication event.

Supplementary Text

Co-expansion patterns of AM-expanded gene families can provide insight into plant regulation of AM symbiosis.

We explored co-expansion patterns of gene families across plant evolution to determine which genetic processes are potentially co-regulated in AM symbiosis and to identify how the genetic mechanisms of AM symbiosis may have diverged in specific plant lineages like grasses. Identifying gene families that grow and shrink together across evolution (i.e., co-expansion) can inform us of which kinds of mechanisms are potentially co-regulated because they are either experiencing similar selective pressures (i.e., they are important for fitness under similar conditions) or the functional outcomes of these gene families are interconnected (e.g., the products of one gene family are substrates for another). Thus, we examined how AM-expanded gene families co-expand across angiosperms using k-means clustering (Figure 2) to explore potentially co-regulated AM-symbiotic processes across angiosperms and in grasses.

The 429 gene families expanded in mycorrhizal-associating plants can be clustered into approximately 6 co-expanding groups that may identify important patterns of molecular evolution. For example, Cluster 1 is one of the most consistently reduced clusters in non-mycorrhizal plants compared to AM plants (W = 819,883.0, FDR = 2.95 × 10^-136^, Wilcoxon Rank-Sum Test, Figure 2), suggesting that the processes encoded by the gene families in Cluster 1 are genetic mechanisms potentially unique to and essential for AM symbiosis. The extensive loss of gene families in Cluster 1 across non-mycorrhizal plants may be due to it containing gene families that exclusively function in AM-symbiosis, such as membrane-bound and arbuscular-localized genes functioning at the symbiotic interface between AM fungi and plant cortical cells. For instance, Cluster 1 contains receptors for fungal-derived enzymes (e.g., EIX2 in gene family OG0000040; *64*, *65*), Vapyrin-like proteins (e.g., gene family OG0012090; Vapyrin is essential to arbuscule formation in AM symbiosis; *66*), pathogenesis/nodulation-associated proteins (e.g., gene family OG0000085), and LysM-containing membrane proteins (e.g., gene family OG0012081). Additionally, Cluster 6 is also highly conserved in AM plants (W = 75,542.5, FDR = 1.28 × 10^-59^, Wilcoxon Rank-Sum Test, Figure 2) and contains several gene families known to regulate AM symbiosis such as the transcription factor RGL1/RAD1 (gene family OG0008696) and CYCLOPS (gene family OG0011811) which links symbiosis-induced changes in calcium signaling to changes in gene expression. Future work determining how these transcriptional regulators regulate AM symbiosis through other members of Cluster 6 would be of interest such as understanding how RGL1 and CYCLOPS may impact regulation of the extracellular and apoplastic space (e.g., gene families OG0001572, OG0013173, and OG0000105) and recognition of signals from symbionts (e.g., gene families OG0009771 and OG0011710). Thus, by leveraging this comparative genomic analysis, we can expand our understanding of even well-characterized genes in AM symbiosis and identify new areas for future research. For instance, our analysis expands our understanding of how even well-characterized genes function by providing evidence for specific mechanisms for how transcriptional regulators, like RGL1 and CYCLOPS, shape AM symbiosis (e.g., linking modifications of the extracellular and apoplastic space to accommodation of mycorrhizal arbuscules by RGL1). We predict that, as more plant species are sequenced and the quality of their assemblies and annotations improve, we can gain even finer resolution of co-expansion patterns across AM symbiosis as well as employ other statistical approaches (e.g., network analyses of expansion patterns) that can provide even further insight into relationships between gene families.

Our clustering approach also suggests that the regulation of AM symbiosis was modified ~65 MYA between the rise of the order Poales and true grasses (i.e., the family Poaceae). We find that, while both Clusters 2 and 3 are expanded across AM plants generally, these gene families diverged in the evolutionary expansion patterns in grasses. Gene families in Cluster 2 shrink/disappear in grasses and those in Cluster 3 expand even more in grasses than in the other AM-plants (W = 517,442.5, FDR = 2.79 × 10^-97^, W = 94,970.0, FDR = 2.09 × 10^-70^, Wilcoxon rank-sum tests, Poaceae vs all other AM plants; Figure 2). This divergence in co-expansion patterns from other AM-associating plants could be an evolutionary signal of a dramatic restructuring of AM symbiosis in grasses (Clusters 2 and 3 represent ~30% of all AM-expanded gene families). For example, symbiosis with relatively ubiquitous symbionts like Epichloë fungi may have taken over functions previously provided by AM fungi thus leading to loss of Cluster 2 gene families and coopting of gene families in Cluster 3 to control both AM fungi and an alternative symbiont. Functionally, Clusters 2 and 3 represent gene families that can regulate the symbiotic interface in grasses with 50.7% (34/67) and 43.5% (27/62) of gene families representing membrane-associated proteins in Clusters 2 and 3, respectively, with Cluster 3 also containing several families that modify the extracellular space/cell wall of plant cells (9.7%). Consequently, these results imply that Clusters 2 and 3 represent substantial reengineering of AM regulation directly at the symbiotic interface and may reflect adaptations to AM symbiosis as a result of unique evolutionary adaptations/pressures in grasses (e.g., modifications to root anatomy). Moving forward, testing how these expanded gene families interact with the unique biology of grasses (e.g., localization of Cluster 3 genes with another symbiont) would provide insight into why AM symbiosis may be alternatively regulated in grasses.

In summary, exploring how gene families expand together across evolution can provide important insight into the underlying mechanisms of host regulation in AM symbiosis both universally and within specific lineages. Our results indicate that regulation at the symbiotic interface is an important component of the evolution of AM symbiosis in plants. Moving forward, as more plant genome annotations reach high-quality, chromosome-scaffold levels, finer evolutionary patterns of co-expansion can be explored (e.g., specific transcription factors co-expanding with specific classes of nutrient transporters). We find indications of finer patterns beyond the 6 clusters in our analysis such as those present/absent in Brassicaceae in Clusters 4 and 5 which may become clearer as more high quality genetic data becomes available for more plants (especially other nonmycorrhizal lineages like Caryophyllales and Cyperaceae). In essence, using co-expansion analyses to identify genes and molecular functions important for AM symbiosis can be useful to holistically expand our understanding of the molecular mechanisms underpinning AM symbiosis by highlighting new genes and molecular processes for future focused, mechanistic studies.

Gene family size and strategies for managing persistent and episodic stress during symbiosis.

Our finding that gene family size can facilitate fine-tuning of context-dependent expression in species interactions creates new questions about how the evolution of context-dependent molecular mechanisms shapes ecological strategies. For example, could larger gene families be more important for fitness under long-term stressors like low phosphorus availability, while smaller gene families provide the advantage of a faster, although cruder, switch in gene expression underlying fitness when a stressor is episodic (e.g., drought)?

Larger gene families create more points of genetic regulation which is useful for having multiple regulatory mechanisms that can help maximize fitness benefits from an interaction. However, having more points of genetic regulation also means that switching between gene expression profiles with vastly different phenotypic outcomes (e.g., to associate and not associate with a partner) may take longer than if there were fewer checkpoints (*67*). This increase in regulatory checkpoints may slow down switching between phenotypic outcomes because more genes and all the associated downstream processes must reach a new equilibrium. For example, we find a positive relationship between gene family size and context-dependent expression under persistent soil chemical stressors like low phosphorus, low potassium, and high salinity stress because larger AM-expanded gene families have more context-dependent expression under these persistent stressors (Figure 3a). This may be due to plants having the benefit of time to follow an ecological strategy that maximizes fitness benefits from associating with AM fungi because natural changes in these environmental factors occur on long time scales (e.g., years; (*68*–*70*). Thus, when long-term environmental factors like phosphorus availability change, larger gene families provide the benefit of fine-tuning gene expression profiles that maximize AM benefits without the negative consequences of plants being too slow to take advantage of environmental changes before their competitors.

In contrast, we find an inverse relationship under drought (i.e., smaller gene families have more context-dependent expression; Figure 3a). This means that complexity is underutilized during drought and suggests that regulation with greater molecular complexity generated by gene family expansions may be disadvantageous under drought stress. Smaller gene families could underlie ecological strategies that prioritize less-fine tuned and more binary responses to dealing with episodic changes in water availability. In fact, the importance of smaller gene families to drought responses has been shown in *Arabidopsis thaliana* through loss-of-function adaptations to drought (i.e., gene loss; *71*) and differential expression analyses in the grass *Pseudoroegneria (Nevski) Löve* demonstrate that genes from contracted gene families are three times more likely to be differentially expressed during drought (*72*). These results, taken together with our finding of greater context-dependent regulation of AM symbiosis in smaller gene families, supports the hypothesis that smaller gene families may be an integral genomic feature of drought responses in general. In contrast to persistent stressors like low nutrient availability, the episodic/shorter-term nature of drought may create a “time crunch” for plants to alter their gene expression profiles quickly so they are not outcompeted by competitors that are more quickly able to adapt to changes in water availability. In essence, the underutilization of genetic complexity from expanded gene families may be the molecular syndrome underlying a different ecological strategy that prioritizes faster responses to episodic stress during symbiosis at the cost of being less fine-tuned than under persistent stress.

In short, whether the relationship between gene family size and context-dependent expression is positive (i.e., larger gene families are more important) or negative (i.e., smaller gene families are more important) may explain the molecular syndromes underpinning alternative ecological strategies. We hypothesize that larger gene families are more important for ecological strategies that prioritize fine-tuning of interactions under long-term stressors and smaller gene families are more important for ecological strategies that prioritize faster responses to species interactions under more episodic stressors. So, while larger gene families may be useful for finding the gene expression profiles that maximize fitness benefits of species interactions when stress occurs on longer time scales, smaller gene families may be more important when the speed of switching between gene expression profiles is more important than fine-tuning gene expression.

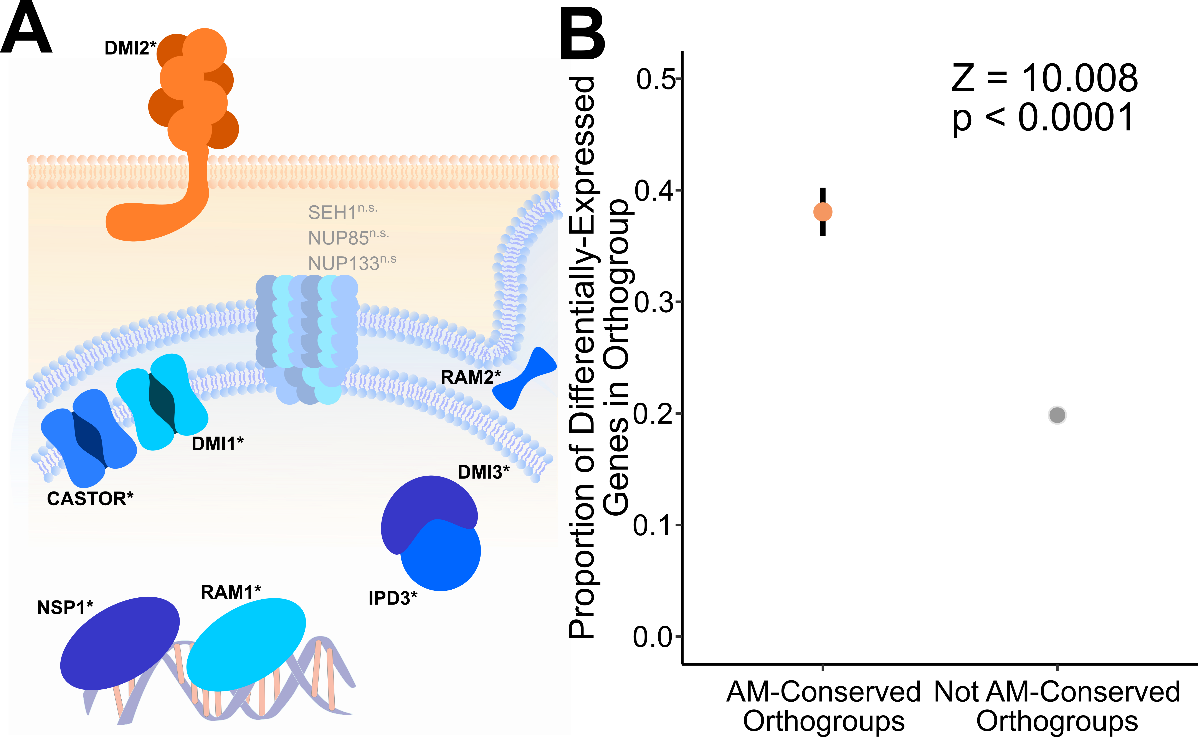

Fig. S1. AM-expanded gene families are involved in AM symbiosis.

(**A**) The Common Symbiosis Pathway (CSP) is an essential pathway in establishing symbiosis. Of the 10 gene families in the CSP, 7 CSP gene families are conserved in AM plants. The three exceptions are the three nuclear pore complex proteins, which may reflect processes that are essential for establishing AM symbiosis but are not unique to it. Unfaded proteins and names reflect the expanded gene families in AM plants (Mann-Whitney U-test, FDR < 0.05). (**B**) Dot-plot of ratio of genes expressed within AM-expanded and non-AM-expanded gene families when plants were grown with and without AM fungi (34, 58). Points represent the mean expression ratio. Error bars represent standard error of the mean. Both points have error bars, but the error bar of non-AM-expanded gene families is small enough to stay within the mean’s point. Z-statistic and p-value are derived from a permutational test. Type or paste caption here. Create a page break and paste in the figure above the caption.

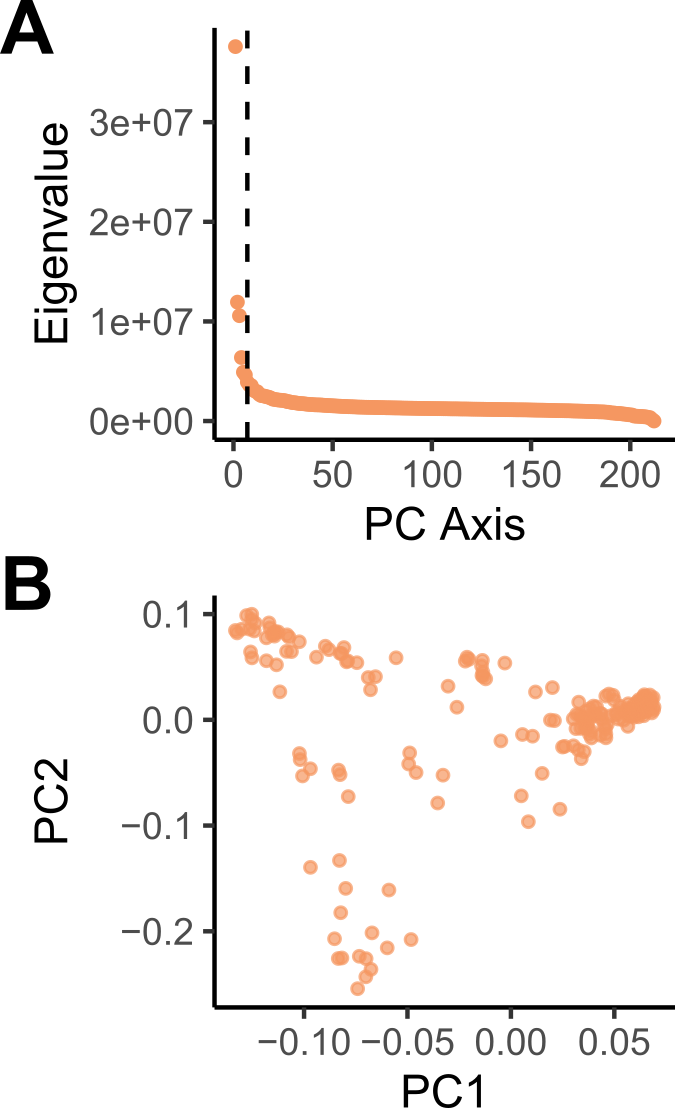

Fig. S2. PCA identifies 6-8 latent factors in *M. truncatula* genotypes.

(**A**) Scree Plot of eigenvalues for PCA of SNP occurrences in 212 M. truncatula populations used in the GWAS spanning the entire Mediterranean (i.e., *M. truncatula*’s range) suggests 6-8 groupings because the distribution of eigenvalues begins to plateau at approximately 6-8 components. Dotted line represents 7 components. (**B**) PCA of SNP occurrences. Visual assessment of the first two axes suggests the existence of 6-8 groupings (i.e., latent factors).

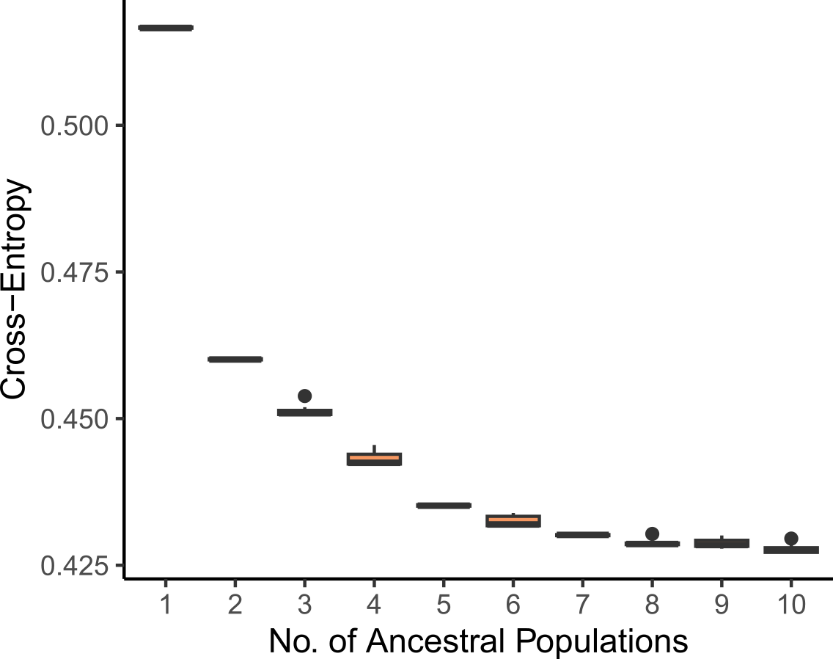

Fig. S3. Cross-entropy analysis of admixture coefficients suggests 6-8 latent factors best describe the underlying genetic structure of M. truncatula populations.

Boxplot of cross-entropy between different runs of admixture coefficient analyses. Points represent outliers. At 6-8 latent factors (K), cross-entropy of admixture coefficients is near its lowest.eate a page break and paste in the figure above the caption.

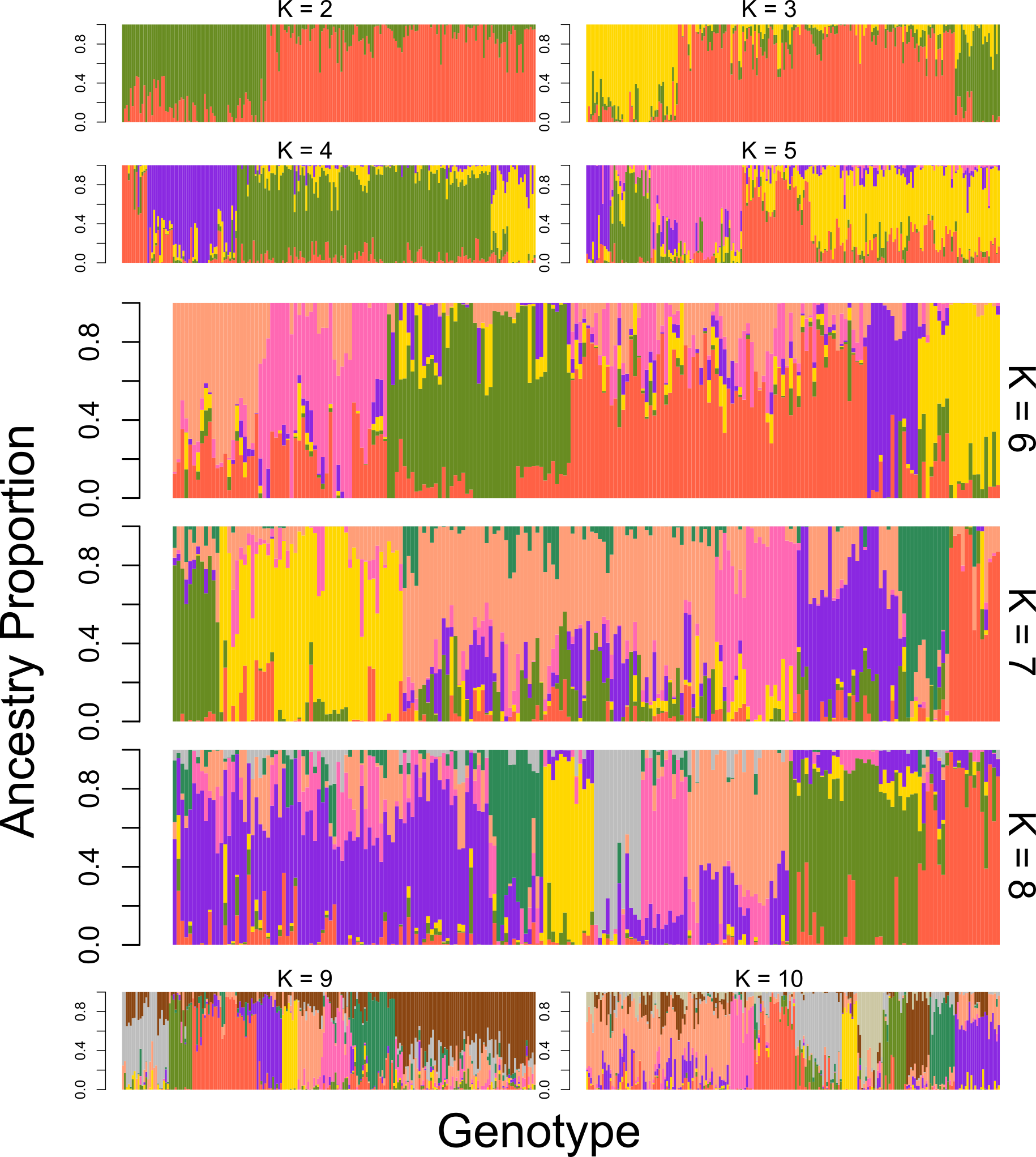

Fig. S4. Visual assessment of Q-matrices also suggests 6-8 latent factors best describe the underlying genetic structure of *M. truncatula*.

Ancestry matrices of the 212 *M. truncatula* populations in the GWAS with 2-10 latent factors (K). Different colors within a plot identify different potential ancestral populations (i.e., latent factors). Colors between plots are not related (i.e., the green group when K=6 is not associated with the green groups in K=7 or K=8).

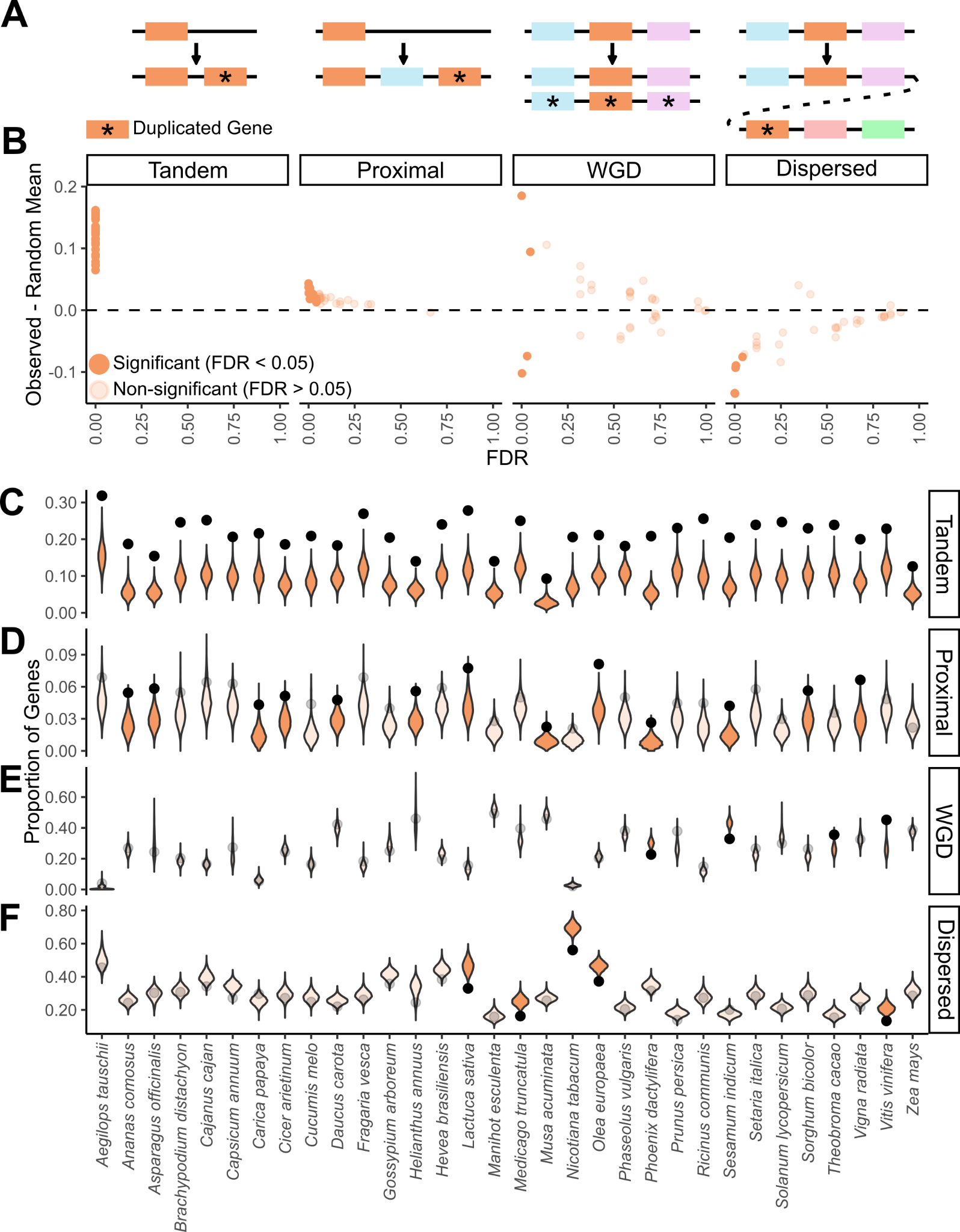

Fig. S5. Tandem duplications are the main driver of growth in AM-expanded gene families across all angiosperm species.

Cartoon schematic of four duplication classes evaluated – tandem, proximal, segmental/whole-genome duplication (WGD), and dispersed duplications – with focal genes in orange and duplicates genes marked by asterisks. **(B)** Frequency of different duplication events in AM-expanded gene families compared to the whole genome across 32 angiosperms. Each point represents one angiosperm species. Y-axis displays the difference between the actual frequency of that type of duplication in AM-expanded gene families and the null expectation (averages from 10,000 randomizations; displayed as violin distributions in C-F). X-axis shows the Benjamini-Hochberg-corrected p-values (FDR) of each comparison between the actual frequency in AM-expanded gene families and the frequencies in the random subsamples. Significant points above the dashed line (dashed line represents no difference between observed and random mean) represent significant enrichment of that duplication type in that species and significant points below the dashed line represent depletion. **(C-F)** Violin plots showing which type of duplications occur significantly more (or less) often in AM-expanded gene families than expected by chance for each of the 32 plant species. Black points represent the proportion of genes in AM-expanded gene families originating from tandem (C), proximal (D), whole-genome/segmental (E), and dispersed (F) duplications found in our analyses. Violin distributions display the null expectations (i.e., distribution of genes that were classified as each duplication type in 10,000 random subsamples of all gene families) for all 32 tested species. Unfaded points and violins are significant relationships (FDR < 0.05; observed proportion of duplications outside of 95% of the null distribution) and faded points and violins are non-significant.

Data S1. (separate file)

Excel file containing metadata and processed data used in statistical analyses.
